# Molecular pathways to high-level azithromycin resistance in *Neisseria gonorrhoeae*

**DOI:** 10.1101/2020.12.02.409193

**Authors:** J.G.E. Laumen, S.S. Manoharan-Basil, E Verhoeven, S Abdellati, I De Baetselier, T Crucitti, B.B. Xavier, S Chapelle, C Lammens, C Van Dijck, S Malhotra-Kumar, C Kenyon

**Author notes:** Address correspondence to Jolein Laumen.

## Abstract

**Objectives:** The prevalence of azithromycin resistance in *Neisseria gonorrhoeae* is increasing in numerous populations worldwide. The aim of this study was to characterize the genetic pathways leading to high-level azithromycin resistance.

**Methods:** A customized morbidostat was used to subject two *N. gonorrhoeae* reference strains (WHO-F and WHO-X) to dynamically sustained azithromycin pressure. We tracked stepwise evolution of resistance by whole genome sequencing.

**Results:** Within 26 days, all cultures evolved high-level azithromycin resistance. Typically, the first step towards resistance was found in transitory mutations in genes *rplD*, *rplV* and *rpmH* (encoding the ribosomal proteins L4, L22 and L34 respectively), followed by mutations in the MtrCDE-encoded efflux pump and the 23S rRNA gene. Low-to high-level resistance was associated with mutations in the ribosomal proteins and MtrCDE-encoded efflux pump. However, high-level resistance was consistently associated with mutations in the 23S ribosomal RNA - mainly the well-known A2059G and C2611T mutations, but also at position A2058G.

**Conclusions:** This study enabled us to track previously reported mutations and identify novel mutations in ribosomal proteins (L4, L22 and L34) that may play a role in the genesis of azithromycin resistance in *N. gonorrhoeae*.

## Introduction

*Neisseria gonorrhoeae* has developed antimicrobial resistance (AMR) to every class of antimicrobials used against it.^1,2^ In order to prevent the further emergence of resistance, treatment guidelines have advised the use of combination therapy with ceftriaxone and azithromycin.^1,3^ Despite this attempt, the prevalence of azithromycin resistance is increasing in Europe and elsewhere.^4,5^ Of even greater concern is the emergence of cases of *N. gonorrhoeae* resistant to both ceftriaxone and azithromycin since 2018.^6^ It is unknown whether combination therapy is retarding or facilitating the emergence of macrolide and cephalosporin AMR.^7^ It is also unclear what role bystander selection, defined as the inadvertent selection of AMR in microorganisms not targeted by antimicrobial therapy, is playing.^8,9^

Azithromycin, a macrolide antibiotic, acts as an inhibitor of protein synthesis by binding within the exit tunnel of the bacterial large ribosomal subunit (50S, consisting of 23S rRNA, 5S rRNA and 31 ribosomal proteins), thereby blocking the exit of nascent polypeptides.^10,11^ Mutations to specific residues in domain V of the 23S rRNA, namely A2059G and C2611T, have previously been reported to confer intermediate to high-level azithromycin resistance.^11–13^

Mutations in the multiple transferable resistance (Mtr) system, are a more common cause of decreased susceptibility to azithromycin.^14^ The Mtr system, a tripartite efflux pump, consists of the periplasmic membrane fusion lipoprotein MtrC and the inner and outer membrane channels MtrD and MtrE.^15–17^ It is responsible for the efflux of a range of antimicrobials, detergents and antimicrobial peptides.^18^ The transcriptional repressor MtrR represses the transcription of the MtrCDE efflux pump operon.^15,19^ Mutations in the MtrR promoter and coding sequences can lead to higher expression of MtrCDE and thereby contribute to macrolide resistance.^20,21^

While mutations in the 23S rRNA azithromycin binding sites and the MtrCDE-encoded efflux pump can explain a large proportion of macrolide resistance, little is known about the pathways leading to macrolide resistance.^22^ The evolutionary steps leading from drug-sensitivity to clinical resistance are usually disentangled using a posteriori comparative genomics of clones isolated from patients before, after and/or during antibiotic treatment.^23^ For *N. gonorrhoeae* these kinds of analyses have revealed several macrolide resistance mechanisms.^24^ A disadvantage of a posteriori comparative genomics is that the clinical strains used for the analyses have already obtained all the geno- and phenotypic changes necessary for resistance. As a consequence, important intermediate adaptive mutations and their response to various selection pressures may be missed.

This provides the motivation to follow the experimental evolution of *N. gonorrhoeae* adapting to azithromycin, as was previously done for ceftriaxone, in real-time and under highly controlled laboratory conditions.^25^ Toprak et al. developed a bioreactor called a morbidostat that allows for continuous culture of bacteria under a constant drug selection pressure based on bacterial growth rate.^26^ Whole genome sequencing is then used to reveal the order, speed and pathways along which resistance evolves. The aim of this study was to investigate the evolution of azithromycin resistance in two *N. gonorrhoeae* WHO reference strains using a customized morbidostat setup.

## Methods

### Strain characteristics

Both WHO reference strains used in this study are susceptible to azithromycin, minimum inhibitory concentrations (MIC) are 0.125 mg/L for WHO-F and 0.25 mg/L for WHO-X.^27^ WHO-F is fully susceptible to ceftriaxone (MIC<0.002 mg/L), whereas strain WHO-X exhibits high-level resistance to ceftriaxone (MIC 2 mg/L).

### Morbidostat setup

The morbidostat was built following the detailed instructions by Toprak et al., modified for *N. gonorrhoeae* according to Verhoeven et al.^26,28^ Of each strain, five morbidostat culture vials, hereafter called cultures, were made. Each culture contained a 10 μL 4.0 McFarland (McF) bacterial cell suspension (WHO-F or WHO-X) in 12 mL gonococcal (GC) broth supplemented with 1% IsoVitaleX (BD BBL™) and vancomycin, colistin, nystatin and trimethoprim selective supplement (VCNT), hereafter called GC medium. The turbidity in each culture was recorded every 60 seconds. After a cycle of 21 minutes, the bacterial growth rate was calculated. When the turbidity was between 1.3-2.0 McF, 1 mL of fresh GC medium was added to stimulate growth. If the turbidity was >2.0 McF and the net growth rate was positive, 1 mL of azithromycin-containing GC medium was added to inhibit growth. As bacteria developed resistance over time, azithromycin concentrations of the GC medium were increased stepwise to regulate growth. After 13 minutes of mixing, 1 mL of excess liquid was removed using a suction pump. The experiment was performed in quintuplet for each strain, with one additional control culture per strain where only GC medium was added throughout the experiment. For further details, see Verhoeven et al.^28^

### Sampling and monitoring of antimicrobial susceptibility

Bacterial suspensions were sampled from each culture two times a week, inoculated onto blood agar and GC-agar plates and incubated for 24 hours at 36.5°C and 6.5% CO_2_. Bacterial cultures from blood agar plates were checked for purity, harvested and transferred in 1 mL of skim milk supplemented with 20% glycerol and stored at −80°C. In addition, bacterial cultures taken from overnight GC-agar plates were suspended in Phosphate Buffered Saline (PBS) to obtain a final turbidity equal to 0.5 McFarland.

The suspensions were plated on GC agar plates, azithromycin E-test gradient strips (bioMerieux, France) were placed and the MICs were determined after overnight incubation.

Once all cultures reached MIC values ≥256 mg/L for azithromycin, the morbidostat experiment was terminated (day 26). Bacterial cultures stored in skim milk supplemented with 20% glycerol were inoculated on blood agar plates. A single colony was inoculated on a new blood agar plate, hereafter called clone, and was used for whole genome sequencing. MICs were determined using agar dilution for ceftriaxone, ciprofloxacin and gentamicin according to the guidelines of CLSI.

### Whole genome sequencing

Genomic DNA was isolated using MasterPure complete DNA and RNA purification kit (Epicentre, USA) according to the manufacturer’s instructions. The DNA concentration was assessed using the Qubit ds DNA HS Assay Kit in a Qubit Fluorometer 3.0 (Thermo Fisher Scientific, USA). Whole genome sequencing of clones was performed via 2×250 bp paired end sequencing (Nextera XT sample) preparation kit and Miseq, Illumina Inc USA.

### Comparative genomic analysis

Reference-based assembly was carried out by mapping reads from the single clones against the references WHO-F (NZ_LT591897) and WHO-X (NZ_LT 592155), with a length fraction of 0.7 and similarity fraction of 0.9. Variants were extracted using the basic variant detection tool implemented in CLC Genomics workbench v20 (clcbio, Denmark).

Azithromycin resistance genes (23S rRNA and *mtrCDE*) and genes that harboured mutations (*rplD*, *rplV*, *rmpH*, *mgtE*, *pilM*, pilin, pre-pilin, *pilQ*, hypothetical proteins, outer membrane beta-barrel protein (OMP), *recJ*, *topB* and *rpoB*) were extracted and manually verified. The allele frequencies of 23S rRNA were determined by deleting the first three of the four 23S rRNA units *in-silico*, followed by reference mapping and variant detection. The proportion of alleles mutated were subsequently calculated as the number of reads mutated divided by the total number of reads covering the position.^29^ Based on the expectation of four genomic 23S rRNA copies in each isolate, the ratios were rounded to the nearest quartile. All the sequences generated in this study were submitted to Bioproject PRJNA664319.

## Results

### *In-vitro* evolution of azithromycin resistance

We performed parallel *in-vitro* experiments for five cultures of strains WHO-F and WHO-X selecting for azithromycin resistance in a modified morbidostat setup. For both WHO-F and WHO-X, the experiment failed in two cultures due to contamination or lack of bacterial growth. In the remaining six cultures, the concentration of azithromycin added to the culture was raised stepwise from 0.5 mg/L medium to 150 mg/L medium for strain WHO-F and from 1 mg/L medium to 300 mg/L medium for WHO-X over the course of the experiment (Figure 1). After 26 days, the MICs of all cultures exceeded 256 mg/L (Figure 1). MICs increased about 2000-fold for WHO-F and 1000-fold for WHO-X, whereas those of the control cultures remained similar to baseline. One culture of WHO-F and all triplicates of WHO-X developed high-level resistance (MIC ≥256 mg/L) between day 13 and 17. High-level azithromycin resistance was attained by day 24 and 26 in the other two cultures of WHO-F (Figure 1).

**Fig. 1.**
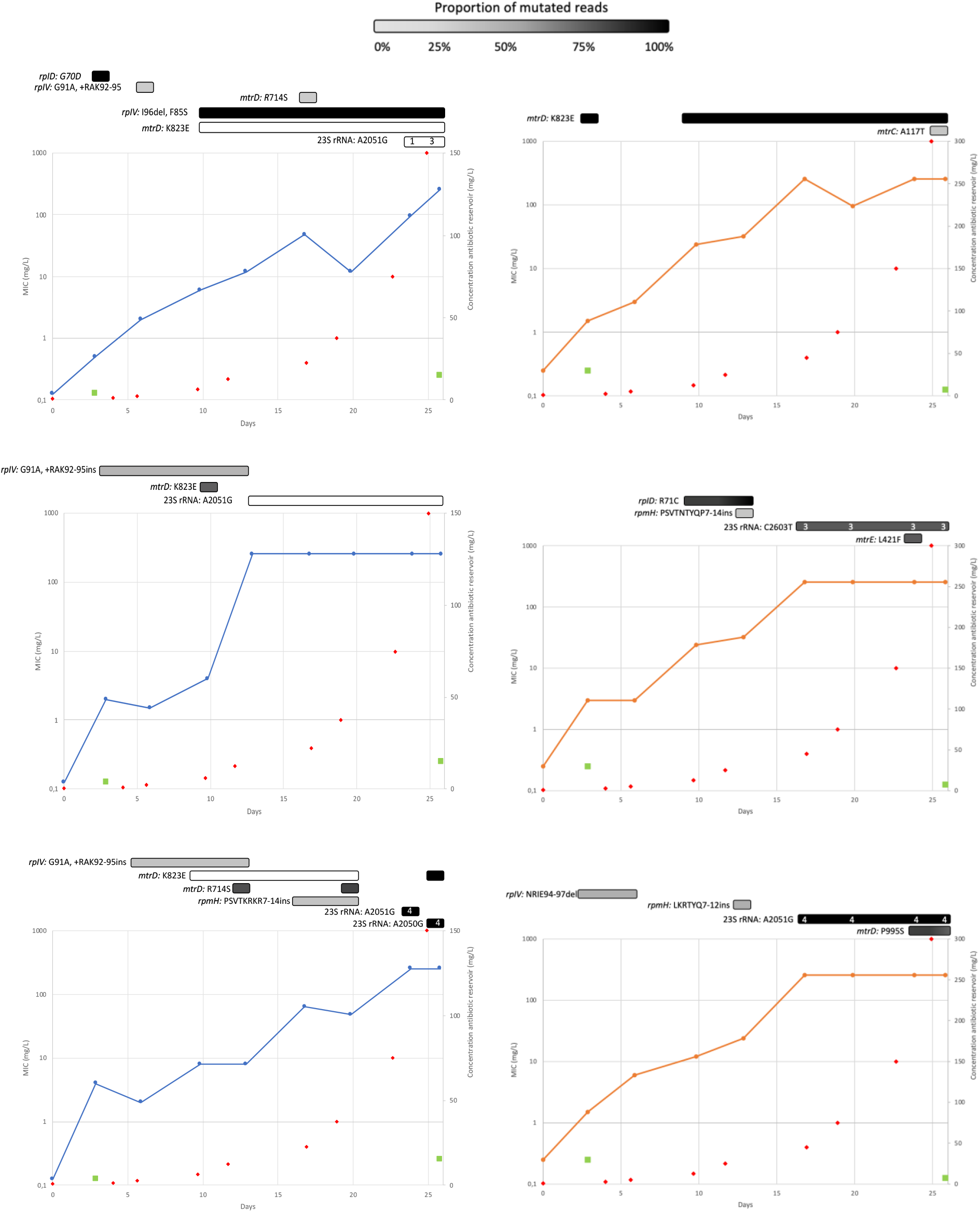
Azithromycin susceptibility of three *N. gonorrhoeae* cultures of WHO-F (left: in blue) and three cultures of WHO-X (right: in orange) over time during a morbidostat experiment. The MICs for azithromycin were tested using E-tests (bioMerieux, France) during the experiment up to a MIC of 256 mg/L. The initial MIC of the WHO reference strains were 0.125 mg/L for WHO-F and 0.25 mg/L for WHO-X. Red squares indicate the azithromycin concentration of the medium added to the experimental vials. Green squares denote the MICs for azithromycin of a control vial, where no azithromycin was added during the experiment. Bars on top represent the mutations found over time, the colour of the bars represent the percentage of reads mutated. For the 23S rRNA gene, the number of alleles mutated are indicated within the bar.

### Allelic variation in 23S rRNA and *mtrCDE*

In all except one culture, all highly resistant clones carried the A2050G (A2058G, *Escherichia coli* numbering), A2051G (A2059G, *E.coli* numbering) or C2603T (C2611T, *E.coli* numbering) single nucleotide polymorphisms (SNPs) that are known to cause macrolide resistance. While the A2050G mutation was present in one clone from strain WHO-F, the A2051G mutation was observed in 12 clones (WHO-F, n=8 and WHO-X, n=4) and the C2603T SNP was present in four clones that evolved from the same culture of strain WHO-X (Table 1). Mutations at all three positions of the 23S rRNA were associated with a MIC of ≥256 mg/L and were acquired between day 13 and 26 (Figure 1). Point mutations in the 23S rRNA and the number of mutated alleles are summarised in Table 1 and Figure 1.

**Table 1.** Mutations in 23S rRNA and *mtrCDE* genes

| Gene | Locus tag |  | Mutation(s) in WHO-F & X cultures by vial number: |  |  |  |  |  |
| --- | --- | --- | --- | --- | --- | --- | --- | --- |
|  | WHO-F | WHO-X | F1 | F2 | F3 | X1 | X2 | X3 |
| 23S<br>rRNA | C7S05_RS07920 | C7S01_RS07170 | A2051G | A2051G | A2050G,<br>A2051G |  | C2603T | A2051G |
|  | C7S05_RS10420 | C7S01_RS09405 |  |  |  |  |  |  |
|  | C7S05_RS11675 | C7S01_RS09915 |  |  |  |  |  |  |
|  | C7S05_RS07155 | C7S01_RS11355 |  |  |  |  |  |  |
| *mtrC* |  | C7S01_RS07540 |  |  |  | A117T |  |  |
| *mtrD* | C7S05_RS08265 | C7S01_RS07535 | K823E,<br>R714S | K823E | K823E,<br>R714S | K823E |  | P995S |
| *mtrE* |  | C7S01_RS07530 |  |  |  |  | L421F |  |

F1, F2 and F3 denote the three culture vials from strain WHO-F. X1, X2 and X3 denote the three culture vials from strain WHO-X. For the 23S rRNA gene, mutations are expressed as SNPs, whereas mutations for other genes are described at amino acid level.

All cultures developed non-synonymous (NS) mutations in genes encoding the MtrCDE efflux pump. The A117T and L421F mutations in genes *mtrC* and *mtrE*, respectively, were found individually in two cultures from the WHO-X strain. Three different NS mutations were found in the *mtrD* gene. The K823E mutation emerged in all three cultures from the WHO-F strain and one culture from the WHO-X strain. Mutations were acquired early on (between day 3-10, MIC 1.5 to 6 mg/L) and remained present while high-level resistance was achieved. Two cultures from strain WHO-F harboured the R714S mutation, present on day 13 and 20 (MIC 8-48). The P995S mutation was only found in one culture from strain WHO-X and was acquired on day 24 (MIC ≥256 mg/L). All mutations are summarized in Table 1.

### Mutations in 50S ribosomal proteins (L4, L22 and L34)

A G70D missense mutation emerged in the L4 protein (*rplD* gene) in one clone of one WHO-F culture. This was associated with low-level resistance (MIC 0.5 mg/L) (Table 2, Figure 1). In one of the WHO-X cultures, the R71C missense mutation was observed in two consecutive clones and was associated with mid-level azithromycin resistance (MIC 24 to 32 mg/L). In total, four and three different mutations were identified in the L22 and L34 ribosomal protein, respectively. Out of the four L22 mutations, three mutations were observed in the WHO-F cultures (n=20 clones) and one in one WHO-X culture (n=2 clones). Mutations in the *rplV* gene were associated with low- to high-level resistance (MIC 1.5 to ≥256 mg/L). Out of the three insertions in the *rpmH* gene, one was found in one WHO-F culture (MIC 48-64 mg/L), whereas the other two mutations were separately found in two WHO-X cultures (MIC 24 and 32 mg/L) (Table 2, Figure 1). Mutations in *rplD* and *rplV* emerged early (day 3 to 10) and mutations in *rmpH* slightly later (day 13 to 17).

**Table 2.** Mutations in ribosomal protein genes (*rplD, rplV* and *rpmH*)

| Gene<br>(protein) | Locus tag |  | Mutation(s) in WHO-F & X cultures by vial number: |  |  |  |  |  |
| --- | --- | --- | --- | --- | --- | --- | --- | --- |
|  | WHO-F | WHO-X | F1 | F2 | F3 | X1 | X2 | X3 |
| *rplD*<br>(L4) | C7S05_RS11295 | C7S01_RS11795 | G70D |  |  |  | R71C |  |
| *rplV*<br>(L22) | C7S05_RS11275 | C7S01_RS11815 | I96del,<br>G91A,<br>+RAKG92-<br>95, F85S | G91A,<br>+RAKG92-<br>95 | G91A,<br>+RAKG92-95 |  |  | NRIE94-97del |
| *rpmH*<br>(L34) | C7S05_RS13510 | C7S01_RS12980 |  |  | PSVTKRKR7-14 |  | PSVTNTYQP7-14 | LKRTYQ7-12 |

F1, F2 and F3 denote the three culture vials for strain WHO-F. X1, X2 and X3 denote the three culture vials for strain WHO-X. Mutations are described at the amino acid level.

### Other variants

Mutations in other genes in which we observed mutant alleles in at least two cultures are summarized in Table 3. The magnesium transporter E (*mgtE*) gene acquired a deletion and three NS mutations in cultures from both WHO strains (MIC 16 to ≥256 mg/L). Three deletions were found in two cultures from WHO-F in the autotransporter OMPs during several timepoints (MIC 4 to ≥256 mg/L). The single-stranded-DNA-specific exonuclease *recJ* acquired the A443V mutation early on and maintained this over time in two cultures from the WHO-F strain (between day 3-26, MIC 1.5 to ≥256 mg/L). A comprehensive list of all mutations is provided in Supplementary Table 1 (Table S1).

**Table 3.** Other mutations repeatedly observed.

| Gene | Locus tag |  | Mutation(s) in WHO-F & X cultures by vial number: |  |  |  |  |  |
| --- | --- | --- | --- | --- | --- | --- | --- | --- |
|  | WHO-F | WHO-X | F1 | F2 | F3 | X1 | X2 | X3 |
| *mgtE* | C7S05_RS07220 | C7S01_RS09475 |  |  | ADV74-76del, R290H |  | E78K, R109Q | ADV74-76del |
| Pre-pilin | C7S05_RS12725 |  |  | D112N, T110A | D67del, K66Q, K63E |  |  |  |
| PilQ | C7S05_RS00605 | C7S01_RS00640 | ins501QEKTE, N501K, del, L437del | R36del | ins509G, R36del, I27del | T708del | V14del |  |
| Hypothetical | C7S05_RS03045<br>C7S05_RS03190 |  | H27del, QFR31-33KQK | H27del, QFR31-33KQK | H27del |  |  |  |
| OMP | C7S05_RS00395<br>C7S05_RS06355<br>C7S05_RS11450 |  | SS13-14Xdel |  | SS20-21delX, LF18-19Xdel |  |  |  |
| *recJ* | C7S05_RS02430 |  |  | A443V | A443V |  |  |  |

F1, F2 and F3 denote the three culture vials for strain WHO-F. X1, X2 and X3 denote the three culture vials for strain WHO-X. Mutations are described at the amino acid level.

### Cross resistance

The MICs for ceftriaxone, ciprofloxacin and gentamicin did not increase over the course of the experiment. Exact values are provided in Supplementary Table 2 (Table S2).

## Discussion

All cultures from both *N. gonorrhoeae* strain WHO-X as well as WHO-F evolved high-level azithromycin resistance in the morbidostat (MIC ≥256 mg/L) within four weeks. Two out of three cultures derived from WHO-F showed stepwise increases in resistance, attaining a MIC of ≥256 mg/L between day 20 and 24. In contrast, all three cultures from strain WHO-X showed a more rapid increase in MIC, attaining a MIC of ≥256 mg/L between day 13 and 17. The difference in the rate at which resistance evolved between WHO-F and WHO-X might be explained by the fact that at baseline WHO-X, but not WHO-F, had a pre-existing single-base-pair (A) deletion in the 13-bp inverted repeat of the MtrR promoter region that is associated with decreased susceptibility to azithromycin.^27^

Overall, mutations tended to occur first in genes encoding the ribosomal proteins (L4, L22 and L34), followed by the MtrCDE-encoded efflux pump and 23S rRNA mutations. Mutations in the *rplD* and *rplV* genes that encode the L4 and L22 ribosomal proteins, respectively, have been associated with macrolide resistance in other organisms such as *E. coli* and *Streptococcus pneumoniae.*^30,31^ Until recently, the role and prevalence of ribosomal protein mutations in conferring macrolide resistance in *N. gonorrhoeae* was unclear. However, a recent bacterial genome-wide association study identified the G70D mutation in the 50S ribosomal protein L4 as being significantly associated with increased azithromycin MICs in clinical isolates of *N. gonorrhoeae*.^32^ The *rplD* G70D mutation was present in 231 out of 4850 isolates (4.76%) and an additional 34 isolates contained other mutations at amino acid positions 68, 69 and 70. These other putative L4 binding site mutations were associated with significantly higher azithromycin MICs compared to both L4 G70D and L4 wild-type strains.

This is in line with our results. We found the G70D mutation in a clone with a MIC of 0.5 mg/L, while the newly observed R71C mutation was seen in two clones with MICs of 24 and 32 mg/L.

The missense mutations in the tentacle of L4 (G70D and R71C) interact with the peptide exit tunnel. Previously, it was proposed that these L4 mutations narrow the peptide exit tunnel so that macrolides are not able to bind to the 50S ribosomal subunit.^33^ Similarly, a deletion of the MKR (positions 82-84) amino acids in ribosomal protein L22 was shown to reduce susceptibility to macrolides via enlarging the loop that forms a constriction in the ribosome exit tunnel. This would in turn allow nascent-chain egress and translation in the presence of bound macrolides.^33^ However, Moore et al. questioned this theory by providing experimental evidence that the MKR deletion in *E. coli* ribosomes reduced erythromycin susceptibility not via blockage of the exit-tunnel but rather by reducing the intracellular erythromycin concentration via increased expression of AcrAB-TolC efflux pumps. This effect was thought to be mediated by increased translation of the relevant mRNAs by reducing programmed ribosome stalling.^34^ Synergistic relationships between L22-mediated antibiotic resistance and efflux systems have also been reported in *Haemophilus influenzae* and *Campylobacter* species.^35,36^ The resistance associated mutations (RAMs) in *rplD* gene could likewise exert their effect either via blockage of the exit tunnel or via influencing the translation of other proteins.^33^

In a similar vein to the findings of Moore et al, *in-vitro* culture experiments with *Mycobacterium smegmatis* have shown that antibiotic-induced mutations in ribosomal proteins result in reduced susceptibility to a range of antimicrobials in a stepping-stone fashion.^38^ Detailed genomic, transcriptomic and proteomic analyses revealed that ciprofloxacin, for example, first selected for mutations in four ribosomal proteins. These mutations resulted in profound alterations in the transcriptome and proteome, which facilitated the acquisition of mutations in other genes that in turn conferred higher lever resistance. Because the ribosomal mutations had a fitness cost, they served as stepping-stones to higher level resistance but were lost once the bacteria had acquired these other mutations.

Although further experimental work is required to establish if ribosomal protein mutations also act as stepping-stones to AMR in *N. gonorrhoeae*, at least three of our findings are supportive of this concept. Firstly, the acquisition of mutations emerged in the following order: first in the ribosomal genes (*rplD, rplV* and *rpmH*) followed by mutations in the MtrCDE efflux pump and the 23s rRNA. Secondly, mutations in *rplD* were transitory and did not persist beyond day 13 in any of the cultures. Thirdly, *rplD/rplV* mutations were followed relatively rapidly by other mutations.

All cultures developed mutations in genes encoding the MtrCDE efflux pump. Except for changes at residues 714 and 823 in the *mtrD* gene, all mutations were found in clones with a MIC ≥256 mg/L. The K823E mutation was acquired early on (MIC 0.75 to 6 mg/L) and retained while high-level resistance was achieved, while the R714S mutation was found in clones with MICs between 8 and 48 mg/L. In a recent study of a clinical *N. gonorrhoeae* isolate possessing a mosaic-like mtr efflux pump locus with reduced susceptibility to antimicrobials, the K823E mutation was found and shown to result in a gain-of-function of MtrD activity suggesting an important functional role for this residue that can contribute to antimicrobial resistance.^39,40^ Cryo-electron microscopic investigations showed that these conserved residues are located within the periplasmic entrance and proximal and distal sites of MtrD and are therefore crucial for the specificity of the pump.^41^

Mutations likely result in enhanced recognition of antimicrobials, leading to increased efflux and hence reduced susceptibility to antimicrobials.

The acquisition of high-level resistance (≥256 mg/L) was linked to mutations in domain V of the 23S rRNA alleles which was observed in all cultures except one. A2059G mutation, a well-described determinant of high-level azithromycin resistance in *N. gonorrhoeae*, ^13,42^was found in four cultures. Remarkably, in one of these cultures, the clone obtained at day 24 harboured the A2059G mutation whilst 2 days later the A2058G mutation was found in all four gene copies. While mutations at this position have been reported to confer macrolide resistance in other bacteria, it has only recently been identified in two high-level azithromycin resistant (MIC ≥256 mg/L) gonococcal isolates from the Gonococcal Isolate Surveillance Project (GISP) study.^43^ In the past, low level azithromycin resistant clinical isolates (MIC 2 to 8 mg/L) were found to contain the C2611T mutation.^12,13^ However, in one of our cultures derived from strain WHO-X, the large increase in MIC from 32 to ≥256 was associated with this mutation in three of the four copies. In this culture, high-level azithromycin resistance may have been the result of a combination of this C2611T mutation following several other mutations including the MtrE part of the efflux pump and transitory mutations in *rplD* and *rpmH*.

Overall, our findings demonstrate the utility of the morbidostat for unravelling pathways towards gonococcal macrolide resistance. Whilst gonococci are very unlikely to be exposed to such constant antimicrobial selection pressure in an *in vivo* setting; they may encounter a diluted version of this scenario in core-group type populations with a high prevalence of gonorrhoea/chlamydia and frequent antibiotic treatments.^44^ A further limitation of our methodology is that there was no opportunity for *N. gonorrhoeae* to acquire AMR via horizontal gene transfer (HGT) from commensal *Neisseria*.

Although HGT has been shown to play an important role in the acquisition of RAMs in the mtrCDE system, it is unknown what role HGT plays in the development of AMR in the ribosomal proteins L4, L22 and L34.^45^ A final limitation is that the sequenced single clones may not represent the total population. However, population sequencing of a subset of samples showed similar results (data not shown). These limitations notwithstanding, the fact that three parallel cultures of two genetically different reference strains revealed commonalities in the pathways towards resistance is instructive.

In conclusion, our results suggest a common order in which the mutations arise: first in the ribosomal proteins, then in the MtrCDE encoded efflux pump and finally within the 23S rRNA gene. Further experimental work is required to assess the extent to which mutations in *rplD, rplV* and *rpmH* act as stepping-stones for subsequent RAMs in other genes. If these mutations are found to be necessary but temporary stepping-stones, then a case could be made to include them in AMR surveillance activities.

## Acknowledgements

We thank Ivo Jansegers and Patrick Nijs (Institute of Tropical Medicine, Antwerp) for their expertise and assistance throughout all the technical aspects of the morbidostat construction.

## Transparency declarations

All authors declare that no conflicts of interest exist. The funders had no role in study design, data collection and interpretation, or the decision to submit the work for publication.

## References

1. Wi T, Lahra MM, Ndowa F, et al. Antimicrobial resistance in Neisseria gonorrhoeae: Global surveillance and a call for international collaborative action. PLoS Med 2017; 14: 1–16.

2. Unemo M, Nicholas RA. Emergence of multidrug-resistant, extensively drug-resistant and untreatable gonorrhea. Future Microbiol 2012; 7: 1401–22.

3. Unemo M. The ‘2012 european guideline on the diagnosis and treatment of gonorrhoea in adults’ recommends dual antimicrobial therapy. Eurosurveillance 2012; 17: 2012.

4. Yasuda M, Ito S, Hatazaki K, Deguchi T. Remarkable increase of Neisseria gonorrhoeae with decreased susceptibility of azithromycin and increase in the failure of azithromycin therapy in male gonococcal urethritis in Sendai in 2015. J Infect Chemother 2016; 22: 841–3.

5. Cole MJ, Spiteri G, Jacobsson S, et al. Overall Low Extended-Spectrum Cephalosporin Resistance but high Azithromycin Resistance in Neisseria gonorrhoeae in 24 European Countries, 2015. BMC Infect Dis 2017; 17: 1–9.

6. European Centre for Disease Prevention and Control. Extensively drug-resistant (XDR) Neisseria gonorrhoeae in the United Kingdom and Australia. ECDC Stock 2018.

7. Unemo M WK. Dual antimicrobial therapy for gonorrhoea: what is the role of azithromycin? Lancet Infect Dis 2018; 18: 486–8.

8. Tedijanto C, Olesen SW, Grad YH, Lipsitch M. Estimating the proportion of bystander selection for antibiotic resistance among potentially pathogenic bacterial flora. Proc Natl Acad Sci U S A 2018; 115: E11988–95.

9. Kenyon C, Buyze J, Spiteri G, Cole MJ, Unemo M. Population-Level Antimicrobial Consumption Is Associated With Decreased Antimicrobial Susceptibility in Neisseria gonorrhoeae in 24 European Countries: An Ecological Analysis. J Infect Dis 2020; 221: 1107–16.

10. Wilson DN. Ribosome-targeting antibiotics and mechanisms of bacterial resistance. Nat Rev Microbiol 2014; 12: 35–48.

11. Unemo M, del Rio C, Shafer WM. Antimicrobial Resistance Expressed by Neisseria gonorrhoeae: A Major Global Public Health Problem in the 21st Century. Microbiol Spectr 2016; 4: 1–32.

12. Ng LK, Martin I, Liu G, Bryden L. Mutation in 23S rRNA associated with macrolide resistance in Neisseria gonorrhoeae. Antimicrob Agents Chemother 2002; 46: 3020–5.

13. Chisholm SA, Dave J, Ison CA. High-level azithromycin resistance occurs in Neisseria gonorrhoeae as a result of a single point mutation in the 23S rRNA genes. Antimicrob Agents Chemother 2010; 54: 3812–6.

14. Harris SR, Cole MJ, Spiteri G, et al. Public health surveillance of multidrug-resistant clones of Neisseria gonorrhoeae in Europe: a genomic survey. Lancet Infect Dis 2018; 18: 758–68.

15. Hagman KE, Pan W, Spratt BG, Balthazar JT, Judd RC, Shafer WM. Resistance of Neisseria gonorrhoeae to antimicrobial hydrophobic agents is modulated by the mtrRCDE efflux system. Microbiology 1995; 141: 611–22.

16. Hagman KE, Lucas CE, Balthazar JT, et al. The MtrD protein of Neisseria gonorrhaeae is a member of the resistance/nodulation/division protein family constituting part of an efflux system. Microbiology 1997.

17. Delahay RM, Robertson BD, Balthazar JT, Shafer WM, Ison CA. Involvement of the gonococcal MtrE protein in the resistance of Neisseria gonorrhoeae to toxic hydrophobic agents. Microbiology 1997; 143: 2127–33.

18. Handing JW, Ragland SA, Bharathan U V., Criss AK. The MtrCDE efflux pump contributes to survival of neisseria gonorrhoeae from human neutrophils and their antimicrobial components. Front Microbiol 2018; 9.

19. Pan W, Spratt BG. Regulation of the permeability of the gonococcal cell envelope by the mtr system. Mol Microbiol 1994; 11: 769–75.

20. Shafer WM. Mosaic drug efflux gene sequences from commensal neisseria can lead to low-level azithromycin resistance expressed by Neisseria gonorrhoeae clinical isolates. MBio 2018; 9: 1–4.

21. Zarantonelli L, Borthagaray G, Lee EH, Shafer WM. Decreased azithromycin susceptibility of Neisseria gonorrhoeae due to mtrR mutations. Antimicrob Agents Chemother 1999; 43: 2468–72.

22. Eyre DW, Silva D De, Cole K, et al. WGS to predict antibiotic MICs for Neisseria gonorrhoeae. J Antimicrob Chemother 2017; 72: 1937–47.

23. Jansen G, Barbosa C, Schulenburg H. Experimental evolution as an efficient tool to dissect adaptive paths to antibiotic resistance. Drug Resist Updat 2013; 16: 96–107.

24. Unemo M, Shafer WM. Antimicrobial resistance in Neisseria gonorrhoeae in the 21st Century: Past, evolution, and future. Clin Microbiol Rev 2014; 27: 587–613.

25. Gong Z, Lai W, Liu M, et al. Novel genes related to ceftriaxone resistance found among ceftriaxone-resistant Neisseria gonorrhoeae strains selected in vitro. Antimicrob Agents Chemother 2016; 60: 2043–51.

26. Toprak E, Veres A, Yildiz S, Pedraza J. Building a morbidostat: an automated continuous-culture device for studying bacterial drug resistance under dynamically sustained drug inhibition. Nat Protoc 2013; 8: 555–67.

27. Unemo M, Golparian D, Sánchez-Busó L, et al. The novel 2016 WHO Neisseria gonorrhoeae reference strains for global quality assurance of laboratory investigations: Phenotypic, genetic and reference genome characterization. J Antimicrob Chemother 2016; 71: 3096–108.

28. Verhoeven E, Abdellati S, Nys P, et al. Construction and optimization of a ‘NG Morbidostat’ − An automated continuous-culture device for studying the pathways towards antibiotic resistance in Neisseria gonorrhoeae. F1000Research 2019; 8: 560.

29. Johnson SR, Grad Y, Abrams AJ, Pettus K, Trees DL. Use of whole-genome sequencing data to analyze 23S rRNA-mediated azithromycin resistance. Int J Antimicrob Agents 2017; 49: 252–4.

30. Gregory ST, Dahlberg AE. Erythromycin resistance mutations in ribosomal proteins L22 and L4 perturb the higher order structure of 23 S ribosomal RNA. J Mol Biol 1999.

31. Tait-Kamradt A, Davies T, Cronan M, Jacobs MR, Appelbaum PC, Sutcliffe J. Mutations in 23S rRNA and ribosomal protein L4 account for resistance in pneumococcal strains selected in vitro by macrolide passage. Antimicrob Agents Chemother 2000.

32. Ma KC, Mortimer TD, Duckett MA, et al. Increased power from bacterial genome-wide association conditional on known effects identifies Neisseria gonorrhoeae macrolide resistance mutations in the 50S ribosomal protein L4. bioRxiv 2020: 2020.03.24.006650.

33. Gabashvili IS, Gregory ST, Valle M, et al. The polypeptide tunnel system in the ribosome and its gating in erythromycin resistance mutants of L4 and L22. Mol Cell 2001; 8: 181–8.

34. Moore SD, Sauer RT. Revisiting the mechanism of macrolide-antibiotic resistance mediated by ribosomal protein L22. Proc Natl Acad Sci U S A 2008; 105: 18261–6.

35. Peric M, Bozdogan B, Galderisi C, Krissinger D, Rager T, Appelbaum PC. Inability of L22 ribosomal protein alteration to increase macrolide MICs in the absence of efflux mechanism in Haemophilus influenzae HMC-S. J Antimicrob Chemother 2004; 54: 393–400.

36. Cagliero C, Mouline C, Cloeckaert A, Payot S. Synergy between efflux pump CmeABC and modifications in ribosomal proteins L4 and L22 in conferring macrolide resistance in Campylobacter jejuni and Campylobacter coli. Antimicrob Agents Chemother 2006; 50: 3893–6.

37. Efflux A, Smeyz P. Mutations in Ribosomal Protein RplA or Treatment with Ribosomal Acting Antibiotics Activates Production of. 2020: 1–5.

38. Gomez JE, Kaufmann-Malaga BB, Wivagg CN, et al. Ribosomal mutations promote the evolution of antibiotic resistance in a multidrug environment. Elife 2017; 6: 1–25.

39. Rouquette-Loughlin CE, Reimche JL, Balthazar JT, et al. Mechanistic basis for decreased antimicrobial susceptibility in a clinical isolate of neisseria gonorrhoeae possessing a mosaic-like mtr efflux pump locus. MBio 2018; 9: 1–15.

40. Chitsaz M, Booth L, Blyth MT, O’mara ML, Brown MH. Multidrug resistance in neisseria gonorrhoeae: Identification of functionally important residues in the mtrd efflux protein. MBio 2019; 10: 1–14.

41. Lyu M, Moseng MA, Reimche JL, et al. Cryo-EM Structures of a Gonococcal Multidrug Efflux Pump Illuminate a Mechanism of Drug Recognition and Resistance. MBio 2020; 11: 1–15.

42. Zhang J, Van Der Veen S. Neisseria gonorrhoeae 23S rRNA A2059G mutation is the only determinant necessary for high-level azithromycin resistance and improves in vivo biological fitness. J Antimicrob Chemother 2019; 74: 407–15.

43. Nash E, Liu H, Schmerer M, et al. P861 Novel mutation conferring high-level azithromycin resistance in Neisseria gonorrhoeae. Sex Transm Infect 2019; 95: A359 LP–A360.

44. Kenyon C. We need to consider collateral damage to resistomes when we decide how frequently to screen for chlamydia/gonorrhoea in preexposure prophylaxis cohorts. Aids 2019; 33: 155–7.

45. Efflux E, Component P, Wadsworth CB. crossm Azithromycin Resistance through Interspecific Acquisition of. 2018; 9: 1–17.

